# Donor-derived mitochondrial DNA variant peptides elicit allo-specific immune response in transplant patients

**DOI:** 10.1101/2020.06.04.134965

**Authors:** Argit Marishta, Yanqin Yang, Xiaomeng Hu, Moon Kyoo Jang, Karen Cuttin, Annette M. Jackson, Helen Luikart, Tobias Deuse, Kiran K. Khush, Sonja Schrepfer, Sean Agbor-Enoh, Hannah Valantine

**Author notes:** Corresponding Author: Hannah Valantine, and Sean Agbor-Enoh.

## Abstract

In stem cell transplant, mitochondrial DNA (mtDNA) nonsynonymous single nucleotide variants (SNVs) between donor and recipient (D-R) trigger alloimmune responses and transplant rejection. Whether similar alloimmune responses occurs in solid-organ transplantation remains unknown, particularly with the presence of human leukocyte antigen mismatches. This study characterized mtDNA SNVs between D-R of 163 human lung transplant pairs, and then, post-transplantation, assessed alloimmune responses against donor-derived mitochondrial peptides using ELISpot to measure interferon gamma (IFNγ) release from recipient’s monocytes. We identified a median of 6 nonsynonymous mtDNA SNVs (Interquartile Range = 4 – 9) per D-R pair. SNVs were predominantly located at *MT-CYB, MT-ATP6*, and *MT-ND3* genes. The number of SNVs was higher in D-R race non-concordant pairs than in race-concordant pairs. Donor-derived mitochondrial peptides triggered a 19.8-fold higher IFNγ release compared to recipient-derived peptide. These findings were validated in heart transplantation and show that donor-derived mitochondrial peptides trigger allo-specific immune responses after transplantation.

## Introduction

Alloimmune response remains a major drive for development of acute and chronic rejection (1). Traditionally, the alloimmune response is thought to be primarily driven by mismatched human leukocyte antigens (HLA). However, acute rejection has been reported in the absence of HLA-directed alloimmune responses(2–5). Mitochondrial-derived protein mismatch between transplant donor and recipient potentially could contribute to alloimmune response. For example, in a stem-cell transplant, where immunologically dominant MHC and other nuclear antigens are matched between the stem cells and recipient, mismatched mitochondria between the stem cells trigger allo-specific immunity (6, 7). Deuse and collaborators utilized somatic cell nuclear transfer to create stem cells with the same nuclear DNA, but with different mtDNA as the host mice. Upon transplantation, host mice developed allo-specific immune responses directed against mitochondrial antigenic mismatch and rejected the stem cells (6). In solid-organ transplantation, particularly heart and lung transplantation, antigenic matching is not performed. Therefore, alloreactivity of mitochondrial antigenic mismatches between donor and recipient deserves investigation, because such mismatches could trigger chronic rejection.

Transplant recipients of non-European ancestry, particularly Black/African American, have higher acute rejection rates across several organ transplant types (8, 9). The reason for higher rejection rates in Black/African American organ transplant recipients is not fully known, and is often attributed to multiple complex immunobiological and socioeconomic factors (10). For example, African American transplant recipients have greater donor HLA variance compared to recipients of European descent (10, 11). Furthermore, the variance of mitochondrial-derived proteins among different human population groups is well established (12). We reasoned that the variability between the donor- and recipient-derived mitochondrial proteins could provide an explanation for higher rejection rates in Black/African American heart transplant recipients. A new understanding of alloimmune response drivers could offer new mechanisms for the prevention of acute and chronic rejection(3, 13, 14). An essential first step towards that understanding would be to characterize mtDNA variability between the donor- and recipient, setting up the rationale for further clinical studies.

Mitochondrial DNA (mtDNA) differs in size and shape from the nuclear genome and is evolutionarily similar to bacterial DNA (15, 16). mtDNA encodes 13 proteins that, in collaboration with other nuclear-encoded proteins, maintain oxidative phosphorylation (OXPHOS) and produce ATP (15, 17). In addition, mtDNA and other mitochondria damage-associated molecular patterns (DAMPs) trigger higher inflammation than nuclear DNA once released in circulation (15). In a transplant setting with genetic mixing, leakage of allograft mitochondrial peptides and DNA possibly sets the stage for inflammation and allo-immune activation (18–21). Therefore, we hypothesize that nonsynonymous mtDNA SNVs result in donor-derived peptides that trigger allo-specific immunity in solid-organ transplant recipients.

This study had three objectives. First, characterize mtDNA SNVs between lung transplant recipients and donors; second, determine if donor-derived peptides with nonsynonymous SNVs trigger allo-specific immune responses in vitro; and third, validate the allo-specific immune response by replicating the analysis in heart transplant patients.

## Methods

### Study design

Patients from two prospective cohort studies, Genomics Transplant Dynamics (GTD: NCT01985412) and Genomics Research Alliance for Transplantation (GRAfT; NCT02423070), were included. GTD is a single center study (Stanford Healthcare) and GRAfT is a multicenter study consisting of five centers in the Washington DC area (INOVA Fairfax Hospital, Medstar Washington Hospital Center, Virginia Commonwealth University Hospital, University of Maryland Medical Center, and Johns Hopkins Hospital). Both studies have similar study designs (22–24). The primary analysis for this study included 163 lung transplant patients, all enrolled while they were on the transplant waitlist. A pre-transplant blood sample was collected from each subject and their transplant donor to isolate mtDNA, complete sequencing, and characterize SNVs. Patients without pre-transplant blood samples were excluded. mtDNA SNVs identified by high-throughput sequencing were validated by two other platforms – digital droplet PCR and targeted sequencing (25). Approximately one-year post-transplantation, we collected recipient peripheral blood mononuclear cells (PBMCs) and used Elispot to determine allo-specific immune responses triggered by donor-derived peptides. To test allo-specific immune responses triggered by donor-derived mitochondrial peptides in other solid--organ transplants, we repeated the analyses in 19 heart transplant recipients. This study was approved by the institutional review boards of the recruiting centers and the National Heart, Lung and Blood Institute. The study design is depicted in Figure 1.

**Figure 1.**
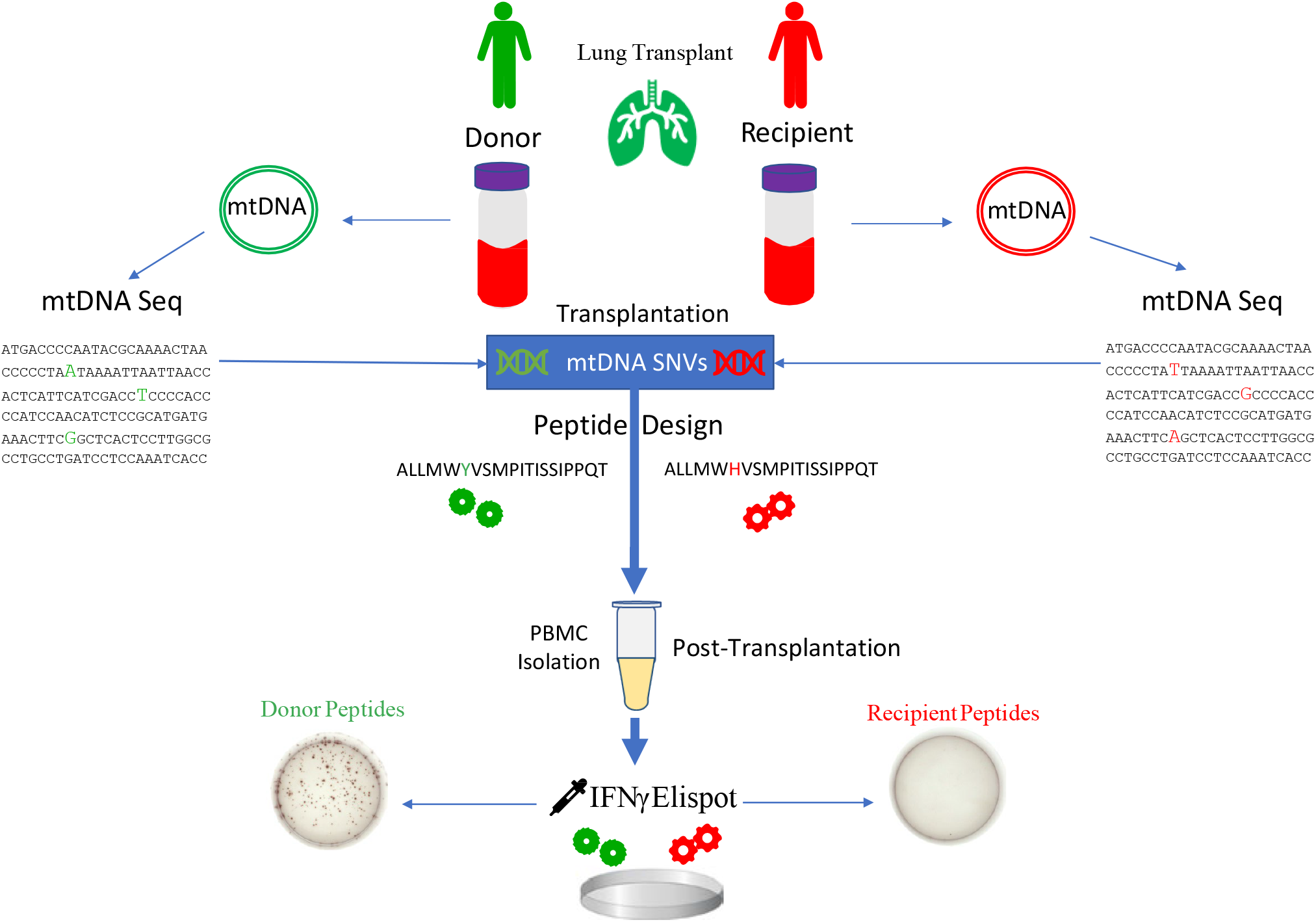
Study design. mtDNA of 163 lung transplant recipients and their organ donors were sequenced. Sequencing reads were analyzed and SNVs for each D-R pair were counted. Peptides at the mitochondria protein coding region differing at one amino acid between the donor and recipient were synthesized. Then, Elispot assay was used to measure the release of IFNγ from peripheral mononuclear blood cells post-transplantation when stimulated with the constructed peptides.

### mt*DNA isolation, library construction and sequencing*

Total DNA, which contains nuclear and mitochondrial DNA, was extracted from whole blood (DNeasy Blood and Tissue kit, Qiagen, Cat No. 69506). Mitochondrial DNA was enriched by step-wise depletion of nuclear DNA. The first step involved incubation of total DNA with proteinA-MBD2-Fc mixture (NEBNext Microbiome DNA Enrichment Kit, New England BioLabs, Microbial enrichment kit, New England Biolabs, Cat No. E2612L), which binds and depletes methylated nuclear DNA fragments. This was followed by exonuclease treatment to digest and deplete linear nuclear DNA strands over circular mtDNA. Quantitative PCR was performed after each step using primers specific to nDNA and mtDNA. mtDNA enrichment efficiency was computed as a ratio of mtDNA-to-nDNA. The enriched mtDNA was then amplified by multiple displacement technology (MDA) using specific primers (REPLI-g Mitochondrial DNA kit, Qiagen, Cat No. 151023), followed by library construction (Nextera XT Sample Prep Kit, Illumina, Cat No. 15032354) and sequencing (pair-end, 50bp) on Illumina HiSeq3000.

### Characterization of SNVs

Sequencing reads were trimmed using Trimmomatic (26) and aligned to the revised Cambridge Reference Sequence (rCRS, accession number: NC_012920) (27) using BWA (v0.7.17) (28). Properly aligned paired-end reads were retrieved and analyzed using samtools (v1.4)(29). SNVs between each D-R pair were called with mpileup2snp in VarScan (v.2.4.3) (30), using the following parameters: minimum-coverage 500, minimum-reads 10, minimum-average-quality 26, minimum-variant-frequency 0.02, minimum-frequency-for-homoplasmy 0.95. The frequency of each single nucleotide variant was computed separately for donor and recipient. Variants were annotated using ANNOVAR database (31) and nonsynonymous SNPs were inputted into UniProtKB database (32) to retrieve peptide sequences.

SNV was present if a D-R pair showed a difference in their variant frequencies of at least 5%. Estimate error rate of sequencing and SNVs calls was 0.1%, so an arbitrary threshold of 5% was selected to reduce erroneous SNVs calls. SNVs were defined as homoplasmic SNV if the variant frequency difference between D-R was greater than 90% or heteroplasmic SNV if the frequency difference was between 5% and 90%.

### Peptide synthesis, PBMC reactivation, Elispot

mtDNA sequences were translated into protein sequences. Nonsynonymous SNVs were defined as single nucleotide variants that lead to mismatched amino acids between D-R. A subset of patients (n=10 patients) were randomly selected. We then constructed and commercially synthesized two 20-mer peptides from each nonsynonymous SNVs position; one with donor amino acid and one with recipient amino acid. Both peptides contained identical amino acid sequences except for a single amino acid difference. One-year post-transplantation, we collected whole blood and isolated PBMCs. We used Elispot to assay for interferon gamma (IFNγ) release as described elsewhere by Deuse and colleagues (25). The number of IFNγ-spots were compared for donor-derived and recipient-derived mt peptides.

### Validation study in heart transplantation

To demonstrate that nonsynonymous donor-derived mtDNA mismatch triggers allo-immunity in other solid--organ transplants, we repeated the analysis in 19 heart D-R pairs. For these patients, we obtained and sequenced donor and recipient pre-transplant mtDNA, characterized the number of SNVs. In a randomly selected subset of patients (n=4), we identified nonsynonymous SNVs, constructed donor-derived and recipient-derived 20-mer peptides and performed Elispot assays. All samples were similarly processed and analyzed as the lung transplant samples.

### Statistical tests

To compare the number of SNVs between demographic groups, we focused the analysis on race and age, given its known association with mtDNA variance (12), and sex, given mitochondria’s maternal inheritance. We employed the non-parametric Wilcoxon signed-rank test to compare the number of SNVs measures between groups, given that the skewness (> 1.05) of the number of SNVs was beyond the limit of normality. Statistical analyses were performed in JMP13 software. For all comparison tests we report median and interquartile range (IQR).

Self-reported sex, age and race were then examined. Each D-R pair was characterized as sex/race concordant (donor and recipient reported the same sex/race) or sex/race non-concordant (donor and recipient reported different sex/race). For age, we used the median age of donors (35 years), and recipients (50 years) to split the cohort into four groups: young donor – young recipient, old donor – young recipient, old donor – old recipient, and young donor – old recipient.

Next, we examined the frequency of SNVs across the different mitochondria genes. To account for the differences in gene length between each of the mitochondria genes, we normalized the analysis for gene length using the following formula: [(gene SNV #) / (mitochondria gene length)] x (16,569 bp mtDNA genome)

## Results

### Description of study cohort

The 163 lung transplant recipients were 51 ± 15 years old at transplantation, 15 years older than their donors (36 ± 14). Half of recipients and two thirds of the donors were males. Among recipients, 76% were White and 12% were African Americans. The majority of donors were White (69%), while African Americans composed 24% of the donor cohort. (Table 1a)

**Table 1a).** Donor and recipient demographics: self-reported sex, self-reported race, and age at transplantation.

| Demographics | Donor |  | Recipient |  |
| --- | --- | --- | --- | --- |
|  | N | Percentage | N | Percentage |
| **Self-Reported Sex** |  |  |  |  |
| F | 66 | 40.49% | 82 | 50.31% |
| M | 9 | 58.28% | 81 | 49.69% |
| Unknown | 2 | 1.23% |  |  |
| **Self-Reported Race** |  |  |  |  |
| Black/African American | 39 | 23.93% | 20 | 12.27% |
| Asian | 7 | 4.29% | 5 | 3.07% |
| Unknown | 5 | 3.07% | 14 | 8.59% |
| White | 112 | 68.71% | 124 | 76.07% |
| **Age at Tx.** |  |  |  |  |
| Mean | 36.24 |  | 51.23 |  |
| Std Dev | 13.88 |  | 15.06 |  |

Race data was unavailable for 29 D-R pairs, leaving 134 D-R pairs with available race data. Race-concordant D-R pairs were 2x more common than race-discordant D-R pairs (n = 93 vs. 41). Similarly, sex-concordant D-R pairs were >2x more common than sex non-concordant D-R pairs (n = 116 vs. 45). D-R pairs were categorized as young donor – young recipient (28%), old donor – young recipient (24%), old donor – old recipient (29%), and young donor – old recipient (19%) (Table 1b).

**Table 1b).**
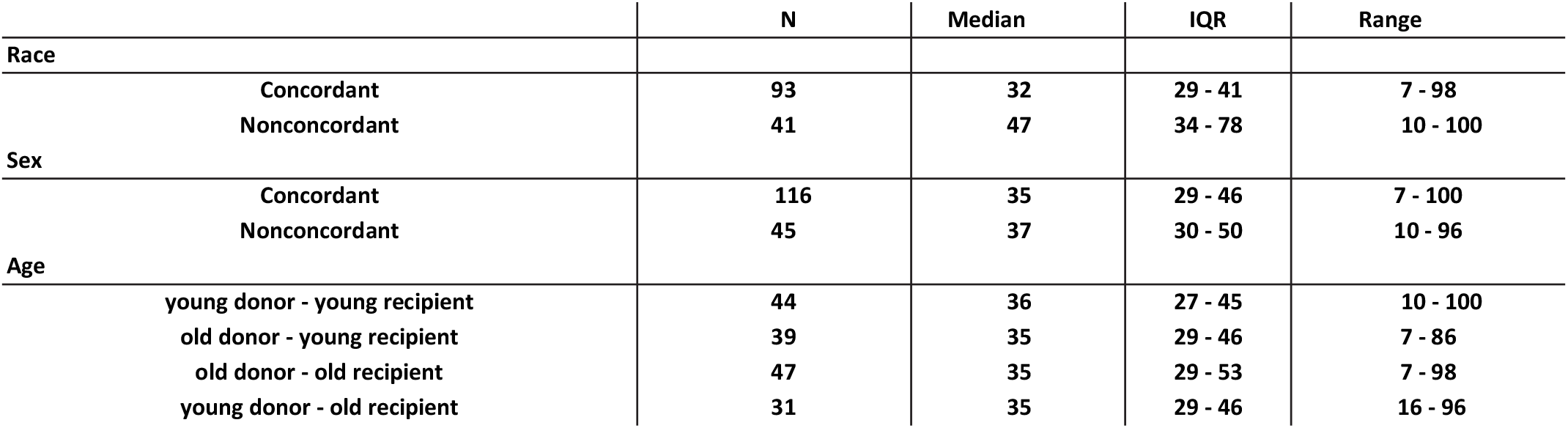
Total homoplasmic SNVs based on groups of donor-recipient race, sex, and age.

|  | N | Median | IQR | Range |
| --- | --- | --- | --- | --- |
| **Race** |  |  |  |  |
| Concordant | 93 | 32 | 29 - 41 | 7 - 98 |
| Nonconcordant | 41 | 47 | 34 - 78 | 10 - 100 |
| **Sex** |  |  |  |  |
| Concordant | 116 | 35 | 29 - 46 | 7 - 100 |
| Nonconcordant | 45 | 37 | 30 - 50 | 10 - 96 |
| **Age** |  |  |  |  |
| young donor - young recipient | 44 | 36 | 27 - 45 | 10 - 100 |
| old donor - young recipient | 39 | 35 | 29 - 46 | 7 - 86 |
| old donor - old recipient | 47 | 35 | 29 - 53 | 7 - 98 |
| young donor - old recipient | 31 | 35 | 29 - 46 | 16 - 96 |

### Characterization of SNVs

Stepwise depletion of nuclear DNA increased mtDNA over nuclear DNA ratio from 10-fold to greater than 10,000-fold. The 326 mtDNAs lung transplant samples (163 donor and 163 recipients) were sequenced to a mean depth of 9540x (Supplementary Table 2.1). We identified a median of 39 SNVs (Interquartile, IQR= 32 – 53) per D-R pair. The majority of the SNVs were homoplasmic with a median of 36 SNVs per D-R pair (IQR = 29 – 46). Heteroplasmic SNVs showed a median frequency of 3 SNVs per D-R pair, (IQR = 2 – 4). A third of all SNVs were in the D Loop (median = 13 SNVs, IQR= 10 – 17 SNVs), leaving two-third of SNVs located in the coding region of mtDNA (median = 27 SNVs, IQR = 21 – 37 SNVs). About a third of the latter SNVs (median = 6 SNVs, IQR = 4 – 9 SNVs) were nonsynonymous non-D-Loop SNVs and lead to proteins with different amino acid sequence between D-R (Table 2). After correcting for gene length, nonsynonymous SNVs were predominantly located in three mt gene regions: *MT-ND3* (0.56 ± 0.62 SNV per D-R, range: 0-3 SNVs per D-R), *MT-CYB* (1.67 ± 0.96 SNVs per D-R, range: 0-4 SNVs per D-R), and *MT-ATP6* (0.93 ± 0.82 SNVs per D-R, range: 0-3 SNVs per D-R). (Supplemental Table 2.2).

**Table 2.** Median (IQR) number of SNVs for the entire lung transplant cohort. SNVs data for the whole mtDNA genome, without the DLoop region, nonsynonymous and without DLoop region, and nonsynonymous protein coding region are shown below. For all these specific regions we calculated the total, homoplasmic, and heteroplasmic SNVs respectively.

| mtDNA SNVs (Lung Tx Cohort) | All |  | Homoplasmic |  | Heteroplasmic |  |
| --- | --- | --- | --- | --- | --- | --- |
|  | Median | IQR | Median | IQR | Median | IQR |
| All SNVs | 39 | 32 - 53 | 36 | 29 - 46 | 3 | 2 - 4 |
| SNVs without DLoop | 27 | 21 - 37 | 25 | 19 - 35 | 1 | 0 - 2 |
| Nonsynonymous SNVs without DLoop | 12 | 9 - 15 | 11 | 8 - 14 | 0 | 0 - 1 |
| Nonsynonymous SNVs (Protein Coding Only) | 6 | 4 - 9 | 6 | 4 - 8 | 0 | 0 - 1 |

### SNVs relationship with self-reported sex, age, and race

The number of SNVs, as well as nonsynonymous SNVs, were similar between sex-concordant and sex non-concordant D-R pairs (median = 11, IQR = 8 −13.75 vs. median 12, IQR = 9 – 14, p = 0.396, Figure 2a). Similarly, all four D-R age groups demonstrated similar amounts of nonsynonymous mtDNA. The young donor-old recipient group showed a trend towards higher number of nonsynonymous mtDNA SNVs compared to old donor-old recipient group, (median = 12 IQR = 11-14 vs median = 10, IQR = 8-14, *p* = 0.067); and to young donor-young recipient, (median = 10 SNVs, IQR = 7-13.75, *p* = 0.077, Figure 2b).

**Figure 2a.**
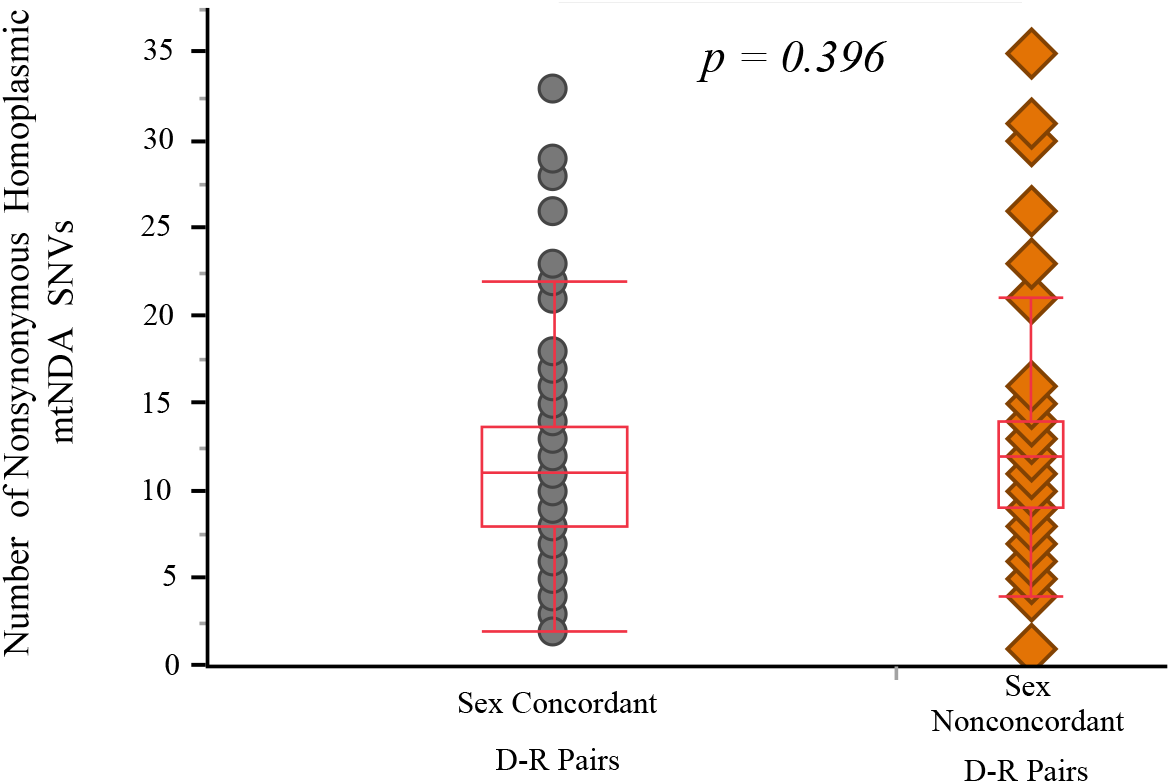
Number of nonsynonymous homoplasmic mtDNA SNVs based on donor-recipient concordant versus nonconcordant sex. The graph shows the number of nonsynonymous homoplasmic mtDNA SNVs for both concordant (grey) and nonconcordant (orange) D-R pairs. Median box plots are shown in red.

**Figure 2b.**
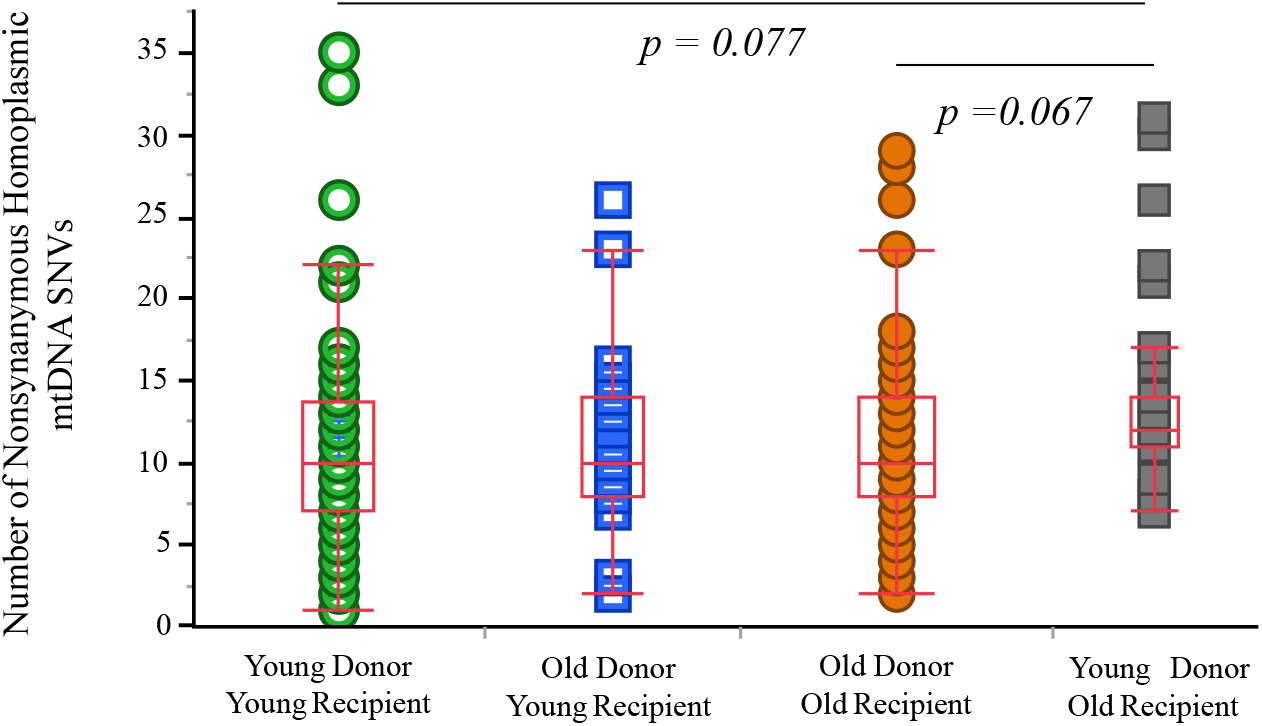
Number of nonsynonymous homoplasmic mtDNA SNVs based on donor-recipient age combinations. We grouped the cohort based on the median age of the recipient and donor into young donor-young recipient, old donor-young recipient, old donor-old recipient, young donor-old recipient. The number of non synonymous homoplasmic mtDNA SNVs for each group is plotted. Groups were compared using Non-parametric Wilcoxon test Two p-values comparing groups are shown. Other p-values were >0.1

In contrast to the other two demographic variables, race concordant D-R pairs showed lower number of nonsynonymous SNVs when compared to the race non-concordant D-R pairs (median = 10; IQR = 8-13 vs. median = 12; IQR = 9.5-23, *p* = 0.024, Figure 2c).

**Figure 2c.**
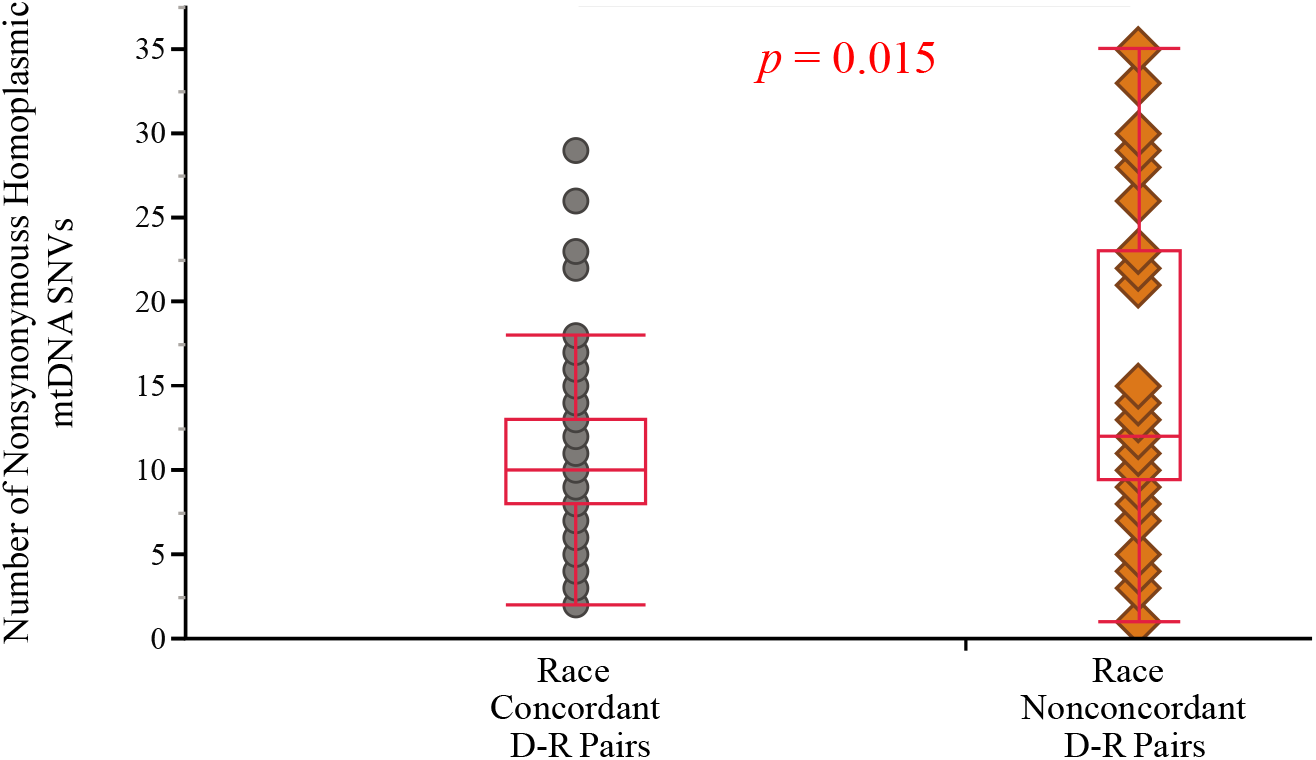
Number of nonsynonymous homoplasmic mtDNA SNVs based on concordant versus nonconcordant donor-recipient self-reported race. The graph shows the number of nonsynonymous homoplasmic mtDNA SNVs for both concordant (grey) and nonconcordant (orange) self-reported race D-R pairs. Median box plots are shown in red.

**Supplementary Table 2.1.** Description of the number of mtDNA reads and mtDNA mapping rate for our sequencing results

|  | Mean (sd) |
| --- | --- |
| Raw Reads | 5088401 (±5381323) |
| Mapping Rate | 81% (± 7%) |
| Avearage Coverage | 38532 (±35950) |
| Trimmed Mapped Reads (Removed Duplicate reads) | 1667481.4 (±962395) |

**Supplementary Table 2.2.** The average nonsynonymous number of SNVs in the lung transplant cohort per mitochondrial gene region.

| Gene Symbol | Average SNVs | Standard Deviation | Minimum SNVs per DR pair | Maximum SNVs per DR pair | Gene Length | mt Genome | Normalized SNVs per gene length |
| --- | --- | --- | --- | --- | --- | --- | --- |
| MT-ATP6 | 0.90 | 0.81 | 0 | 3 | 681 | 16569 | 21.79 |
| MT-ATP8 | 0.15 | 0.39 | 0 | 2 | 207 | 16569 | 11.79 |
| MT-CO1 | 0.37 | 0.82 | 0 | 4 | 1542 | 16569 | 3.96 |
| MT-CO2 | 0.22 | 0.43 | 0 | 2 | 684 | 16569 | 5.35 |
| MT-CO3 | 0.23 | 0.45 | 0 | 2 | 784 | 16569 | 4.80 |
| MT-CYB | 1.55 | 0.93 | 0 | 4 | 1141 | 16569 | 22.45 |
| MT-ND1 | 0.64 | 0.77 | 0 | 5 | 956 | 16569 | 11.06 |
| MT-ND2 | 0.64 | 0.87 | 0 | 4 | 1042 | 16569 | 10.24 |
| MT-ND3 | 0.55 | 0.61 | 0 | 3 | 346 | 16569 | 26.44 |
| MT-ND4 | 0.17 | 0.40 | 0 | 2 | 1378 | 16569 | 1.99 |
| MT-ND4L | 0.02 | 0.15 | 0 | 1 | 297 | 16569 | 1.37 |
| MT-ND5 | 1.04 | 1.34 | 0 | 6 | 1812 | 16569 | 9.54 |
| MT-ND6 | 0.25 | 0.49 | 0 | 2 | 525 | 16569 | 7.94 |
| MT-RNR1 | 1.36 | 1.37 | 0 | 6 | 954 | 16569 | 23.65 |
| MT-RNR2 | 2.41 | 1.57 | 0 | 7 | 1559 | 16569 | 25.62 |
| MT-TA | 0.06 | 0.24 | 0 | 1 | 69 | 16569 | 14.73 |
| MT-TC | 0.04 | 0.20 | 0 | 1 | 66 | 16569 | 10.78 |
| MT-TD | 0.24 | 0.43 | 0 | 1 | 68 | 16569 | 58.30 |
| MT-TE | 0.03 | 0.17 | 0 | 1 | 69 | 16569 | 7.37 |
| MT-TF | 0.02 | 0.15 | 0 | 1 | 71 | 16569 | 5.73 |
| MT-TG | 0.06 | 0.24 | 0 | 1 | 68 | 16569 | 14.95 |
| MT-TH | 0.02 | 0.15 | 0 | 1 | 69 | 16569 | 5.89 |
| MT-TI | 0.01 | 0.11 | 0 | 1 | 69 | 16569 | 2.95 |
| MT-TK | 0.02 | 0.13 | 0 | 1 | 70 | 16569 | 4.36 |
| MT-TL1 | 0.01 | 0.11 | 0 | 1 | 75 | 16569 | 2.71 |
| MT-TL2 | 0.28 | 0.45 | 0 | 1 | 71 | 16569 | 64.43 |
| MT-TM | 0.04 | 0.20 | 0 | 1 | 68 | 16569 | 10.46 |
| MT-TN | 0.01 | 0.08 | 0 | 1 | 73 | 16569 | 1.39 |
| MT-TP | 0.01 | 0.08 | 0 | 1 | 68 | 16569 | 1.49 |
| MT-TQ | 0.08 | 0.27 | 0 | 1 | 72 | 16569 | 18.35 |
| MT-TR | 0.14 | 0.35 | 0 | 1 | 65 | 16569 | 35.97 |
| MT-TS1 | 0.06 | 0.23 | 0 | 1 | 69 | 16569 | 13.26 |
| MT-TS2 | 0.05 | 0.22 | 0 | 1 | 59 | 16569 | 13.78 |
| MT-TT | 0.38 | 0.56 | 0 | 2 | 66 | 16569 | 95.49 |
| MT-TV | 0.02 | 0.13 | 0 | 1 | 69 | 16569 | 4.42 |
| MT-TW | 0.03 | 0.17 | 0 | 1 | 68 | 16569 | 7.47 |

Note: These numbers are arbitrary units.

### Allo-specific immunity from nonsynonymous SNVs

Donor-derived peptides demonstrated 26-fold higher IFNγ release compared to recipient-derived 20mer peptides, p < 0.001 (recipient-peptides: median spots number = 2.3, IQR(1.4 – 5.9); donor-peptides: median spots number=57.5, IQR(39.3 - 109.8); Figure 3). Individual IFNγ release plots for the 10 lung transplant subjects studied is displayed in Supplementary Figure 3.1. Moreover, IFNγ release was when comparing PBMCs collected at one-year post-transplant to PBMCs collected beyond one year after transplantation (Supplementary Figure 3.2).

**Figure 3.**
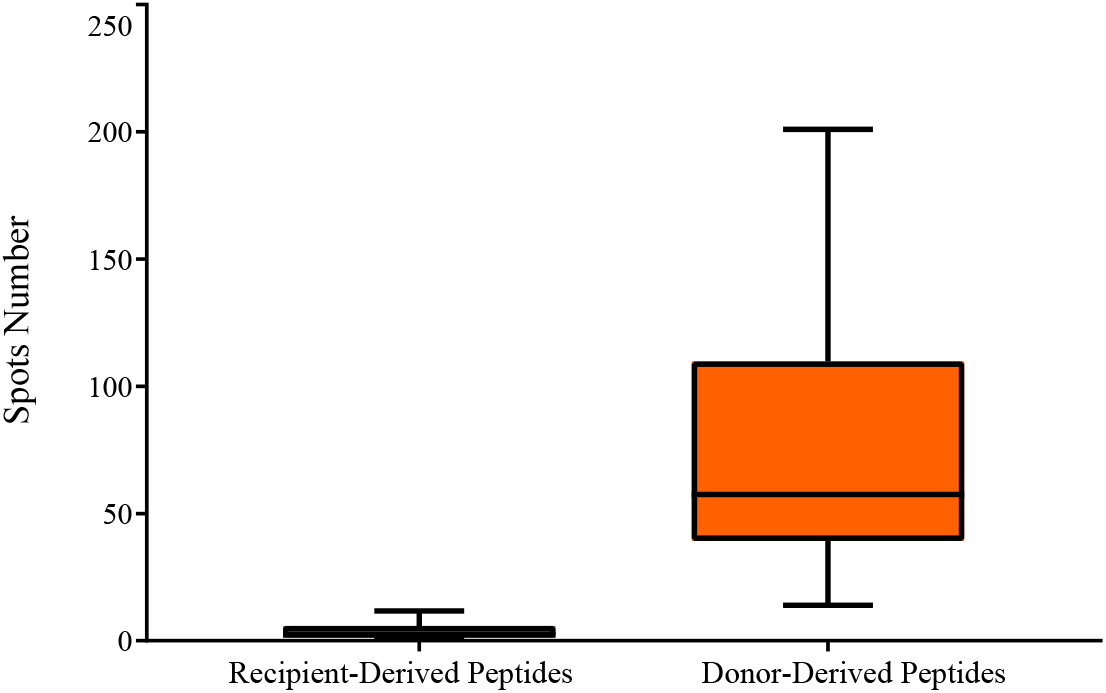
The median number of IFNγ spots for the lung transplant patients. The box plots represent the number of spots measured for each peptide: the orange box plot represent the measurements from the donor-derived peptides and the gray box plot represent the response measured from the recipient-derived peptides. For each individual spot the average of quadruplicate measurement was used. Each individual peptide’s response is shown in the supplementary figure 1.

### Validation of the study in a Heart Transplant Cohort

To validate our findings, we sequenced the mtDNA for 19 heart transplant D-R pairs. SNVs were similar in the heart and lung transplant cohort. We identified a median of 48 SNVs (IQR= 38-80) per D-R pair. The majority of the SNVs were homoplasmic (median = 43 SNVs, IQR= 3-74), with few heteroplasmic SNVs (median = 3, IQR = 2-5). A third of all SNVs were in the D Loop (median = 15, IQR = 12-19), leaving two-third of SNVs located in the coding region of mtDNA (median = 33, IQR = 24-61). About a quarter of the latter SNVs (median = 8, IQR = 6-12) were nonsynonymous non-D-Loop SNVs and lead to proteins with different amino acids sequence between D-R (Table 3). Nonsynonymous SNVs were predominantly located in three mitochondrial gene regions: *MT-ND3* (0.63±0.48 SNV per D-R, range: 0-3 SNVS per DR), *MT-CYB* (1.32±0.97 SNVs per DR, range: 0-4 SNVs per DR), and *MT-ATP6* (1.16±0.67 SNVs per DR, range: 0-3 SNVs per DR), (Supplementary Table 3.1).

**Table 3.**
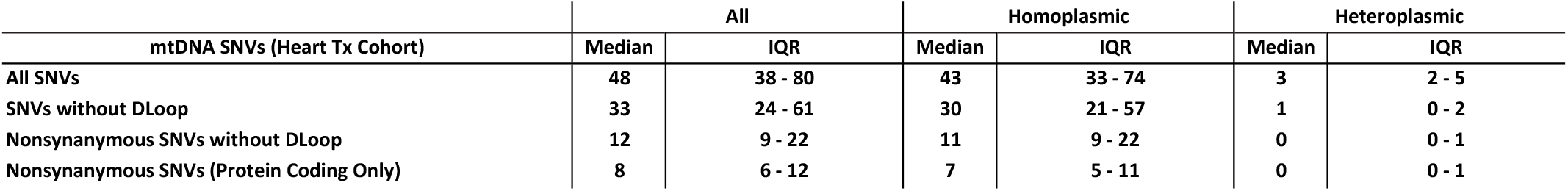
Median (IQR) number of SNVs for the validation heart transplant cohort. SNVs data for the whole mtDNA genome, without the DLoop region, nonsynonymous and without DLoop region, and nonsynonymous protein coding region are shown below. For all these specific regions we calculated the total, homoplasmic, and heteroplasmic SNVs respectively.

Donor-derived peptides demonstrated 47-fold higher IFNγ release compared to recipient-derived 20mer peptides, p < 0.001 (recipient-peptides: median spots number=1.6, IQR(0.8 – 2.2); donor-peptides: median spots number=75.8, IQR(38.1 - 148.1); Figure 4). The individual IFNγ release spot number for each of the 4 heart transplant subjects studied is displayed in Supplementary Figure 4.1. Moreover, IFNγ release was similar when comparing PBMCs collected at one-year post-transplant to PBMCs collected beyond one year after transplantation (Supplementary Figure 4.2).

**Figure 4.**
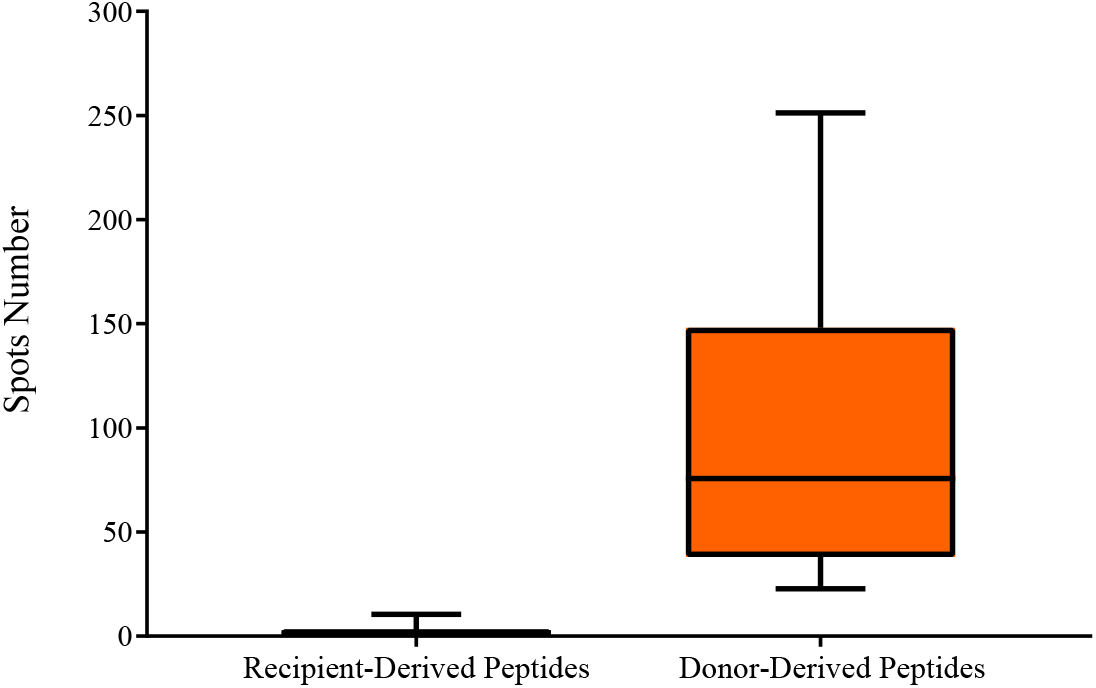
The median number of IFNγ spots for the heart transplant patients. The box plots represent the number of spots measured for each peptide: the orange box plot represent the measurements from the donor-derived peptides and the gray box plot represent the response measured from the recipient-derived peptides. For each individual spot the average of quadruplicate measurement was used. Each individual peptide’s response is shown in the supplementary figure 2.

**Supplementary Figure 3.1.**
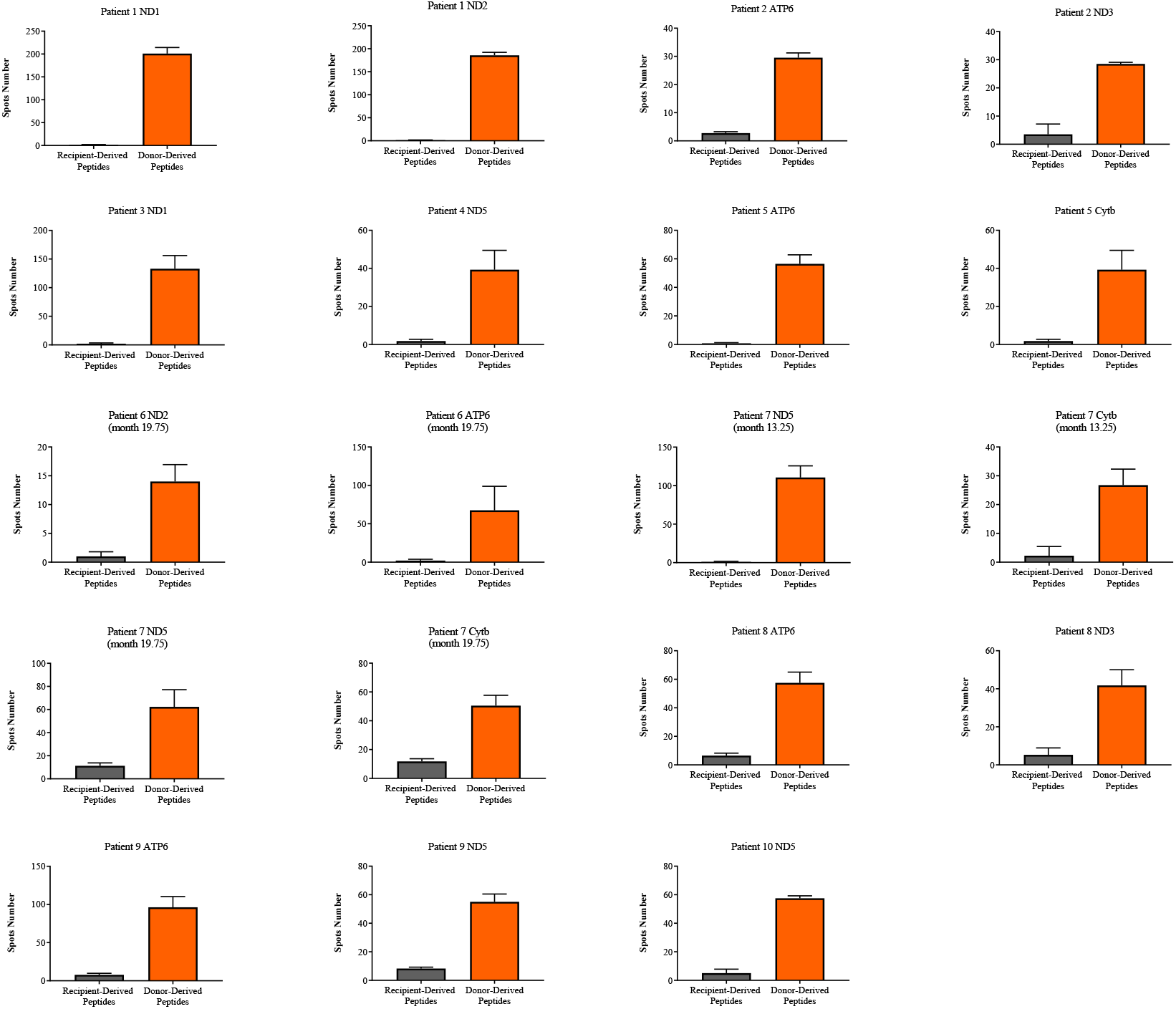
Number of spots measured in Elispot assays for each lung transplant patient. Histograms bars represent the number of spots measured for the donor-derived (gray) and recipient-derived (black) peptides in the Elispot assays. The black lines represent the standard deviation of quadruplicates.

**Supplementary Figure 3.2.**
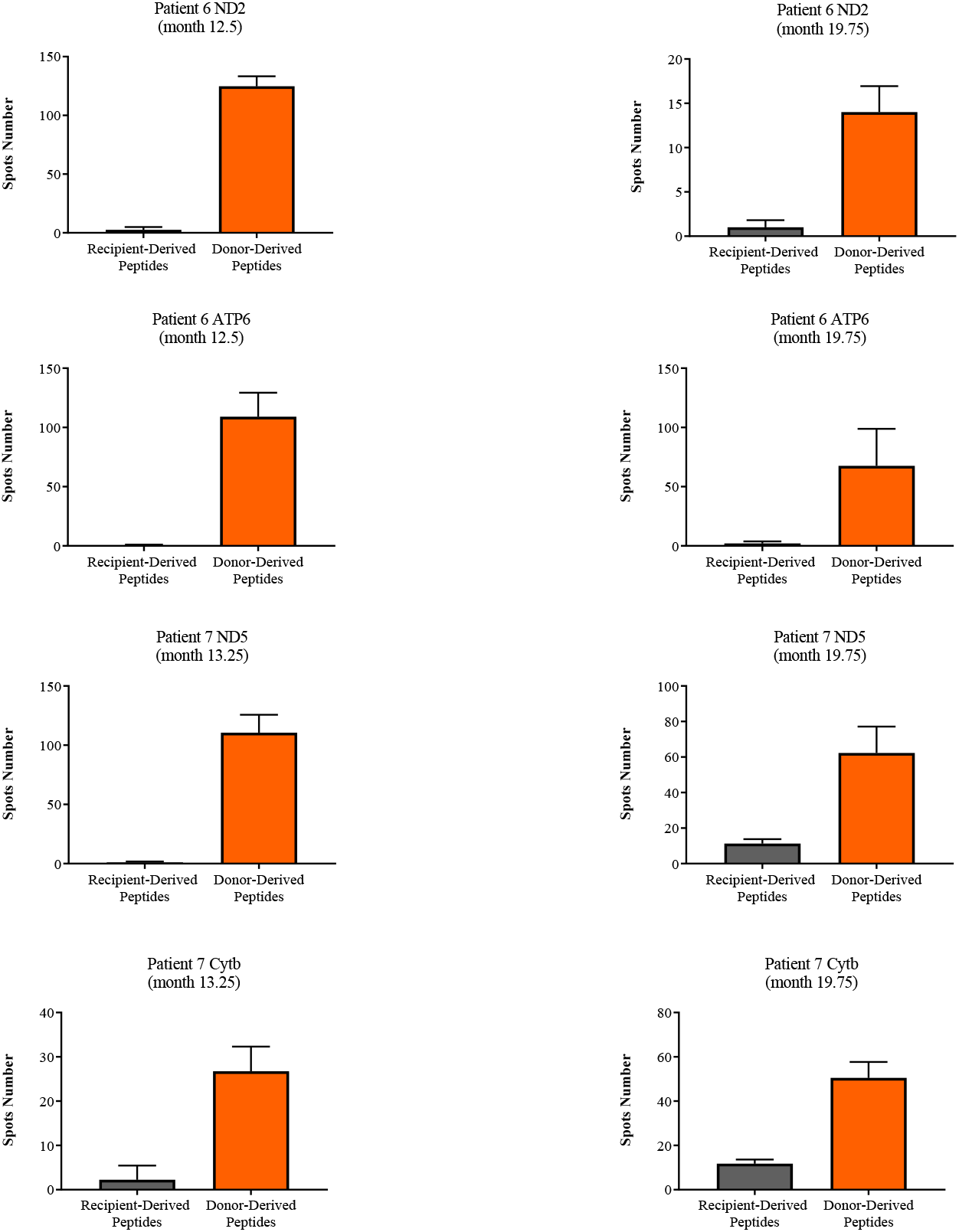
Number of spots measured in Elispot assays for each lung transplant patient at two different PBMC collection timepoints. Histograms bars represent the number of spots measured for the donor-derived (gray) and recipient-derived (black) peptides in the Elispot assays. The black lines represent the standard deviation of quadruplicates.

**Supplementary Table 3.1.** The average nonsynonymous number of SNVs in the heart transplant cohort per mitochondrial gene region.

| Gene Symbol | Average SNVs | Standard Deviation | Minimum SNVs per DR pair | Maximum SNVs per DR pair | Gene Length | mt Genome | Normalized SNVs per gene length |
| --- | --- | --- | --- | --- | --- | --- | --- |
| MT-ATP6 | 1.16 | 0.67 | 0 | 2 | 681 | 16569 | 28.17 |
| MT-ATP8 | 0.16 | 0.36 | 0 | 1 | 207 | 16569 | 12.64 |
| MT-CO1 | 0.63 | 0.81 | 0 | 2 | 1542 | 16569 | 6.79 |
| MT-CO2 | 0.21 | 0.41 | 0 | 1 | 684 | 16569 | 5.10 |
| MT-CO3 | 0.26 | 0.44 | 0 | 1 | 784 | 16569 | 5.56 |
| MT-CYB | 1.32 | 0.98 | 0 | 3 | 1141 | 16569 | 19.11 |
| MT-ND1 | 0.37 | 0.58 | 0 | 2 | 956 | 16569 | 6.39 |
| MT-ND2 | 0.63 | 0.74 | 0 | 2 | 1042 | 16569 | 10.04 |
| MT-ND3 | 0.63 | 0.48 | 0 | 1 | 346 | 16569 | 30.24 |
| MT-ND4 | 0.32 | 0.57 | 0 | 2 | 1378 | 16569 | 3.80 |
| MT-ND5 | 1.63 | 1.22 | 0 | 4 | 1812 | 16569 | 14.92 |
| MT-ND6 | 0.26 | 0.55 | 0 | 2 | 525 | 16569 | 8.31 |
| MT-RNR1 | 1.68 | 1.75 | 0 | 6 | 954 | 16569 | 29.25 |
| MT-RNR2 | 2.74 | 1.77 | 0 | 7 | 1559 | 16569 | 29.09 |
| MT-TA | 0.16 | 0.36 | 0 | 1 | 69 | 16569 | 37.92 |
| MT-TC | 0.11 | 0.31 | 0 | 1 | 66 | 16569 | 26.43 |
| MT-TD | 0.32 | 0.46 | 0 | 1 | 68 | 16569 | 76.95 |
| MT-TG | 0.16 | 0.36 | 0 | 1 | 68 | 16569 | 38.47 |
| MT-TI | 0.11 | 0.31 | 0 | 1 | 69 | 16569 | 25.28 |
| MT-TL1 | 0.11 | 0.31 | 0 | 1 | 75 | 16569 | 23.25 |
| MT-TL2 | 0.16 | 0.36 | 0 | 1 | 71 | 16569 | 36.85 |
| MT-TN | 0.05 | 0.22 | 0 | 1 | 73 | 16569 | 11.95 |
| MT-TP | 0.05 | 0.22 | 0 | 1 | 68 | 16569 | 12.82 |
| MT-TR | 0.16 | 0.36 | 0 | 1 | 65 | 16569 | 40.25 |
| MT-TS1 | 0.05 | 0.22 | 0 | 1 | 69 | 16569 | 12.64 |
| MT-TS2 | 0.05 | 0.22 | 0 | 1 | 59 | 16569 | 14.78 |
| MT-TT | 0.47 | 0.68 | 0 | 2 | 66 | 16569 | 118.92 |
| MT-TW | 0.05 | 0.22 | 0 | 1 | 68 | 16569 | 12.82 |

Note: These numbers are arbitrary units. They do not represent SNVs.

**Supplementary Figure 4.1.**
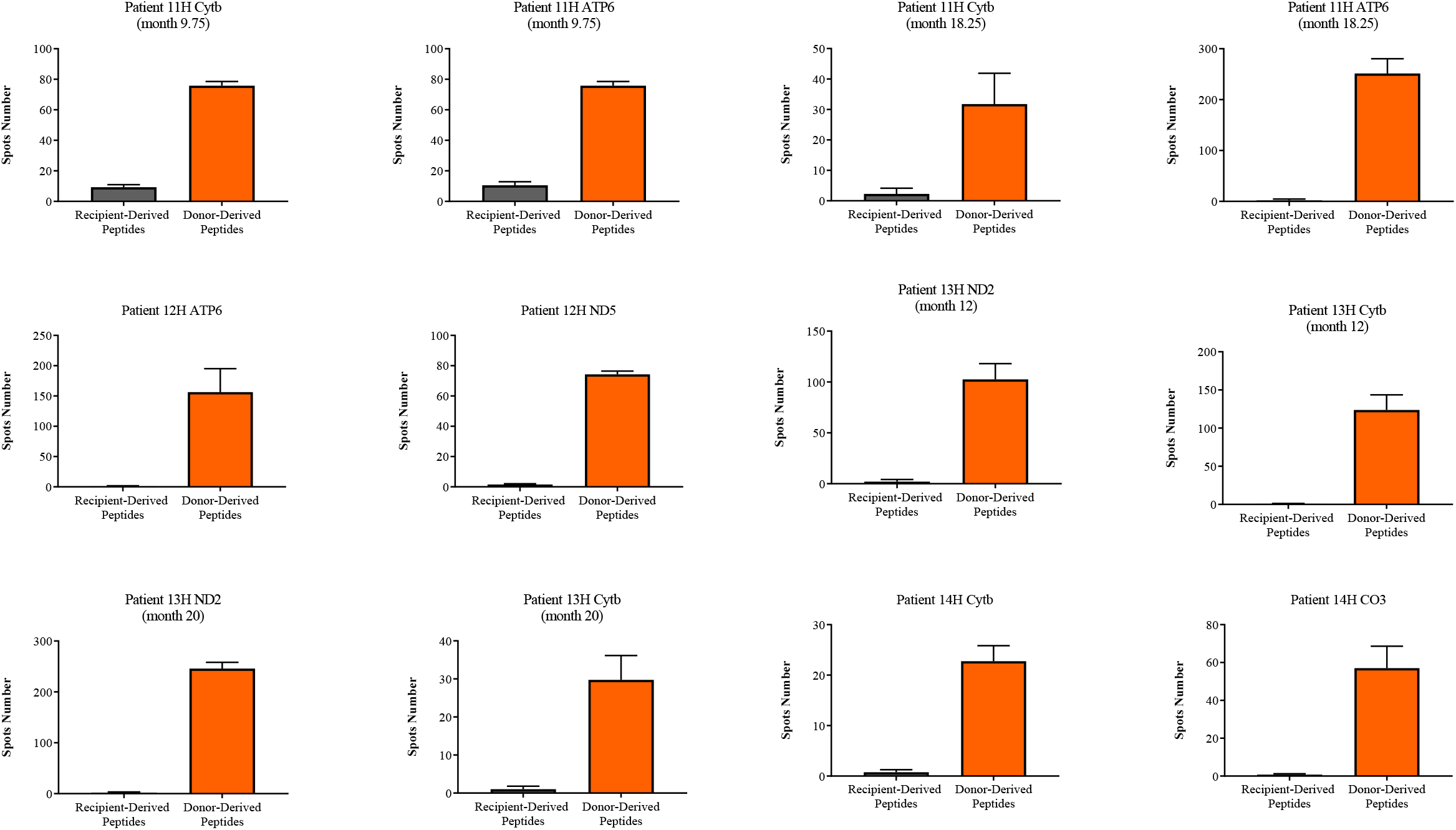
Number of spots measured in Elispot assays for each heart transplant patient. Histograms bars represent the number of spots measured for the donor-derived (gray) and recipient-derived (black) peptides in the Elispot assays. The black lines represent the standard deviation of quadruplicates.

**Supplementary Figure 4.2.**
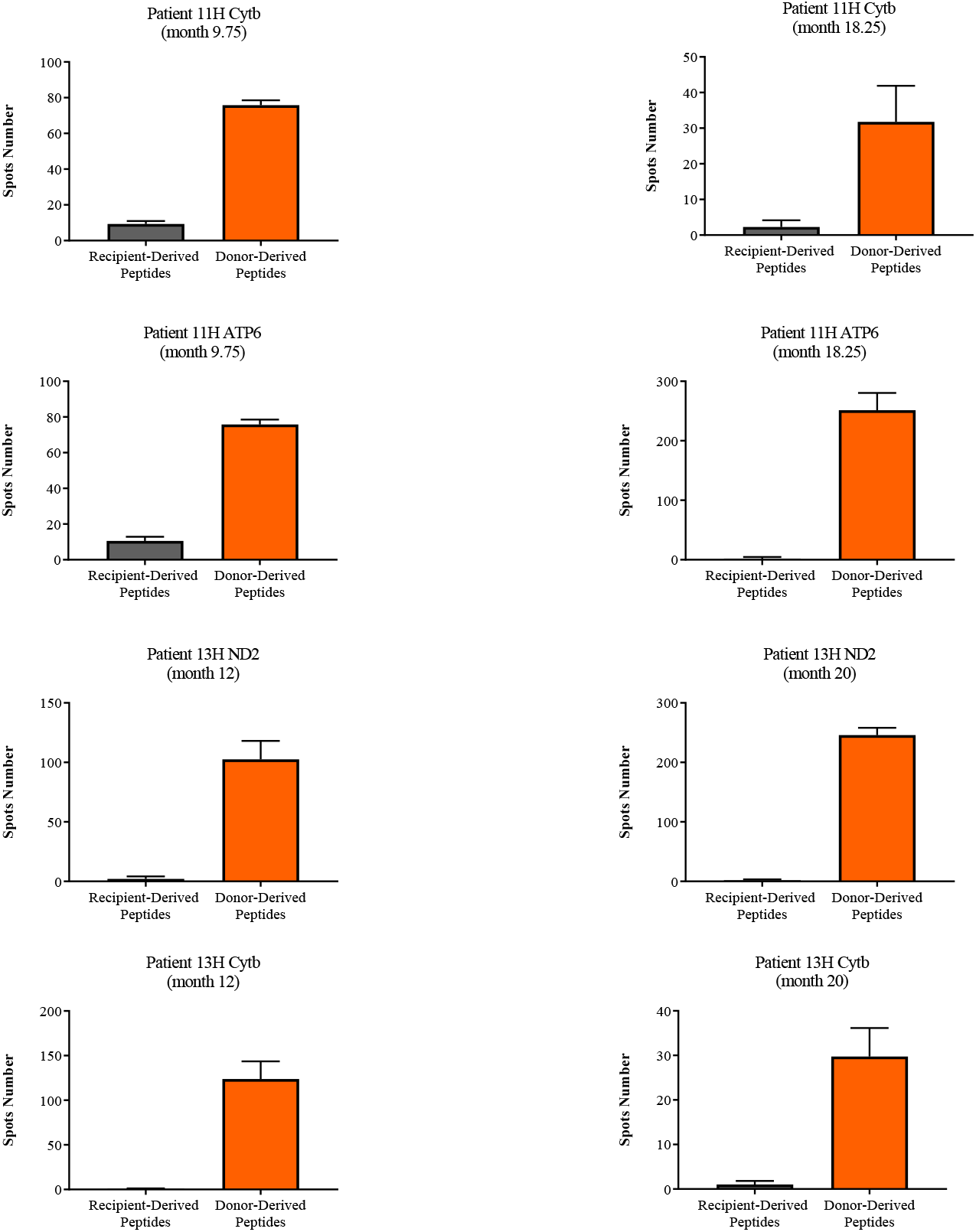
Number of spots measured in Elispot assays for each heart transplant patient at two different PBMC collection timepoints. Histograms bars represent the number of spots measured for the donor-derived (gray) and recipient-derived (black) peptides in the Elispot assays. The black lines represent the standard deviation of quadruplicates.

## Discussion

Release of mitochondrial content exposes transplant recipients to donor-derived proteins and mtDNA -- a very potent DAMP (33, 34). Scozzi and colleagues, as well as others (35), have described the association of circulating mtDNA with severe primary graft dysfunction after lung transplantation and proposed that the graft injury is coupled with neutrophil trafficking (36). We demonstrate that mtDNA peptide variants elicit allo-specific immune responses, which may potentially contribute to allograft rejection. In the absence of rejection, immune responses directed against the mitochondria may orchestrate deleterious effects on the allograft -- given the important cellular function of the mitochondrial, including apoptosis, metabolism, and immune regulation (15, 37). Additionally, cellular stress and necrosis, both common in solid-organ transplantation, promote extracellular mtDNA release (33, 38). Therefore, pharmacologically blocking the extracellular release of mitochondrial components could enhance immunosuppression, prevent allo-immunity, and potentially prevent rejection.

Cyclosporine A, and other drugs, inhibit the release of mitochondrial components through membrane pores(39–41). However, cyclosporine A and other calcineurin antagonists have been part of immunosuppression treatment for decades, but the blocking of mitochondrial membrane pores may not be sufficient to prevent mtDNA and DAMPs release. Other mechanisms of mitochondrial content release are more complex and involve binding of mt components to cellular cargo proteins (42, 43) or extracellular vesicles (44). Determining the ability of other drugs to target these more complex mechanisms would provide an additional avenue for immunosuppression.

In this study, we have shown that transplantation exposes patients to donor-derived SNVs outside the classical MHC region or other nuclear proteins. After transplantation, the donor-derived peptides triggered allo-specific immune responses, measured as IFN-γ release. These findings were consistent in heart and lung transplant patients, suggesting that donor-derived proteins trigger allo-specific immune responses across solid-organ transplant procedures. In addition, demographic factors that contribute to mtDNA diversity, such as race, contributes to the number of SNVs. Race non-concordant D-R pairs demonstrated a higher number of SNVs compared to race-concordant D-R pairs. The implications of these findings are potentially broad and deserve further investigation.

Immunosuppressive therapies and post-transplant monitoring have reduced, but not eliminated, the incidence of acute rejection (1, 45). However, allo-specific immune responses directed to immunodominant antigens, such as HLA, do not explain all cases of rejection (4). Recently, several publications have implicated alloimmune responses directed to non-HLA antigens as causes of rejection. Our findings demonstrate that donor-derived mitochondrial proteins are immunogenic, even in a background of immunodominant HLA and non-HLA nuclear antigenic differences between donors and recipients. Considering the idea of personalized immunosuppression in donor-recipient pairs with low HLA mismatch at molecular level (46), our data suggest that the contribution of mtDNA SNVs, and possibly other known non-HLA factors, to the alloimmune response needs to be included in the personalized immunosuppression plans. The interaction of mitochondrial-directed alloimmune response and alloimmune response directed against immunodominant HLA deserves further investigation. Such investigations should also assess the role of mitochondrial allo-specific immune responses in transplant rejection.

The higher number of SNVs in race non-concordant transplant donor and recipient compared to race-concordant transplants may be associated with poorer outcomes. (10, 47–49). Greater HLA mismatch in race non-concordant transplants is a potential contributor (10, 50). Our work suggests that the higher number of mtDNA SNVs, in combination with genomic SNVs, in race non-concordant donor-recipient pairs may contribute to increased alloimmune responses and poorer outcomes.

Our study had several limitations that should be addressed in future studies. Our analyses focused on allospecific mononuclear cell responses triggered by mitochondrial components and did not investigate concurrent innate immune or antibody-immune responses. We also collected a limited amount of PBMCs, and as such only a subset of nonsynonymous SNVs identified per donor-recipient pair were tested with our Elispot assay. However, despite the small subset of nonsynonymous SNVs tested we observed a significant response over self in all patients tested. Characterization of immune responses directed against each SNV, as well as the cumulative effect of all SNVs for each patient, deserves further investigation. Lastly, we do not assess the relationship of mitochondrial-directed allo-specific immune responses to transplant outcomes such as acute rejection or long-term allograft survival. Our future studies will collect PBMCs at pre-specified time-points to enable such analyses.

In conclusion, we demonstrate that transplantation exposes patients to donor-derived mitochondrial SNVs and proteins. Some of these donor proteins have amino acid differences, compared to recipient proteins, that trigger allo-specific immune responses. The role of the observed allo-specific immune responses in transplant rejection, and the interaction of such immune responses with other known HLA and non-HLA directed responses deserves further investigation.

